# Rapid and Efficient Ambient Temperature X-ray Crystal Structure Determination at Turkish Light Source

**DOI:** 10.1101/2022.10.12.511637

**Authors:** Mehmet Gul, Esra Ayan, Ebru Destan, J Austin Johnson, Alaleh Shafiei, Abdullah Kepceoğlu, Merve Yilmaz, Fatma Betül Ertem, İlkin Yapici, Bilge Tosun, Nilüfer Baldir, Nurettin Tokay, Zeliş Nergiz, Gözde Karakadioğlu, Seyide Seda Paydos, Cahine Kulakman, Cengiz Kaan Ferah, Ömür Güven, Necati Atalay, Enver Kamil Akcan, Haluk Cetinok, Nazlı Eylül Arslan, Kardelen Şabanoğlu, Bengisu Aşci, Serra Tavli, Helin Gümüsboğa, Sevde Altuntaş, Masami Otsuka, Mikako Fujita, Şaban Tekin, Halilibrahim Çiftçi, Serdar Durdaği, Ezgi Karaca, Burcu Kaplan Türköz, Burak Veli Kabasakal, Ahmet Kati, Hasan DeMirci

## Abstract

High-resolution biomacromolecular structure determination is essential to better understand protein function and dynamics. Serial crystallography is an emerging structural biology technique which has fundamental limitations due to either sample volume requirements or immediate access to the competitive X-ray beamtime. Obtaining a high volume of well-diffracting, sufficient-size crystals while mitigating radiation damage remains a critical bottleneck of serial crystallography. As an alternative, we introduce the plate-reader module adapted for using a 72-well Terasaki plate for biomacromolecule structure determination at a convenience of a home X-ray source. We also present the first ambient temperature lysozyme structure determined at the Turkish Light Source (*Turkish DeLight*). The complete dataset was collected in 18.5 mins with resolution extending to 2.39 Å and 100% completeness. Combined with our previous cryogenic structure (PDB ID: 7Y6A), the ambient temperature structure provides invaluable information about the structural dynamics of the lysozyme. *Turkish DeLight* provides robust and rapid ambient temperature biomacromolecular structure determination with limited radiation damage.

## INTRODUCTION

X-ray crystallography has played a dominant role in understanding the structural dynamics of biomacromolecules and elucidating molecular mechanisms of many important biological processes in the past five decades ^1^. Conventional single crystal X-ray crystallography has led to many scientific developments and discoveries in basic science and medicine and is still considered a relevant structural biology technique to many biologists ^2,3^. However, this approach can result in radiation damage within proteins due to primary X-ray absorption during diffraction data collection ^4,5^. The X-ray photons cause Auger decay and K-shell photoionization and may generate reactive oxygen species that can propagate throughout the crystal ^4,5^. This damage results in the reduction of diffraction data quality and can lead to compositional and conformational structural perturbations ^6^. To overcome this, data collection can be performed at cryogenic temperatures; however, cryogenic data collection does not eliminate radiation damage completely but can itself perturb crystal lattice and protein structures ^7^. In addition, global radiation damage can perturb diffraction patterns and increase the unit-cell volume and mosaicity. The increased unit-cell volume results in nonisomorphism, leading to difficulties in structure determination ^8^.

Cryogenic temperature diffraction data collection allows improved resolution by protecting crystals from radiation damage caused by powerful X-ray sources. However, they result in altered structural conformations of the side chains and loop regions that can potentially deviate significantly from those obtained at near physiological temperature ^9^. Temperature can induce pH changes ^10^ and the addition of cryo-protectants can lead to structural artifacts in cryogenic structures ^11^. These may alter the native structure of the protein and its interactions with ligands or other protein partners within the crystal lattice. Unlike cryo-crystallography, *in situ* data collection at ambient temperature may provide us with invaluable macromolecular structural dynamics information in near-physiological conditions ^12,13^.

Serial femtosecond crystallography (SFX) techniques performed at X-ray free electron lasers (XFELs) can overcome the experimental limitations of conventional X-ray cryocrystallography by mitigating radiation damage through the use of ultra-short femtosecond X-ray pulses ^14,15^. In addition, this technique is more suited for understanding structural dynamics since data collection is performed at ambient temperature. Unfortunately, crystal samples in SFX are consumed in a single-use, making SFX techniques even more challenging than conventional cryo X-ray crystallography ^16^. Therefore, there is a need for groundbreaking, easy-to-use, easy-to-access, and highly-efficient state-of-the-art developments in this field to obtain routine high-resolution crystal structures at ambient temperature. Here we provide a paradigm changing example of a high-resolution protein crystal structure obtained from a home X-ray source *“Turkish DeLight”* at near-physiological temperature by switching to a *“Warm Turkish DeLight”* mode ^17^.

In this study, we introduce a high throughput fully-automated *in situ* single crystal X-ray crystallography data collection technique by using the Rigaku Oxford Diffraction *XtaLAB Synergy-S* diffractometer. We modified the commercial *XtalCheck-S* plate reader system to allow diffraction data collection from low-cost Terasaki crystallization plates. Comparison of cryogenic and ambient temperature lysozyme structures generated using *Turkish DeLight* shows that the *XtalCheck-S* module offers rapid and high-quality data collection in a short period of time. Lysozyme, a structurally well characterized protein, was used to obtain structural insights into differences between cryogenic and ambient structures. The main purpose of this experimental setup is to “*serially*” collect preliminary diffraction data from protein crystals at ambient temperature using a *multiwell*-*multicrystal* plate reader as an alternative to serial femtosecond and millisecond X-ray crystallography (SFX/SMX) techniques performed at XFELs and synchrotrons respectively.

## MATERIALS AND METHODS

### Protein sample preparation and crystallization

Chicken egg lysozyme (Calzyme Laboratories, Inc, USA) was dissolved in nanopure water to a final concentration of 30 mg/mL. The lysozyme protein solution was filtered by a 0.22 μm hydrophilic polyethersulfone (PES) membrane filter (Cat#SLGP033NS, Merck Millipore, USA). The filtered sample was stored in 1.0 mL aliquots at −45 °C until crystallization experiments were performed. Sitting drop vapor diffusion microbatch under oil technique was used for crystallization with approximately 3000 commercial sparse matrix and grid screen crystallization conditions ^17^. Equal volumes of crystallization conditions were mixed with 0.83 μL of 30 mg/mL lysozyme solution (1:1 v/v) in a 72-well Terasaki plate (Cat#654180, Greiner Bio-One, Austria). Then, each well was covered with 16.6 μL of paraffin oil (Cat#ZS. 100510.5000, ZAG Kimya, Turkey) and incubated at 4 °C. Lysozyme crystallized in most crystallization conditions within 24 hours. A compound light microscope was used to observe crystal formation in wells of Terasaki plates.

### Sample delivery and *XtalCheck-S* setup for data collection

Rigaku’s XtaLAB Synergy Flow XRD system controlled by *CrysAlisPro* 1.171.42.59a software (Rigaku Oxford Diffraction, 2022) was used for data collection as described in Atalay *et al*. (2022) ^17^. As opposed to the initial published work, the airflow temperature of Oxford Cryosystems’s Cryostream 800 Plus was adjusted to 300K (26.85°C) and kept constant for data collection at ambient temperature. Instead of the intelligent goniometer head (IGH), the 72-well Terasaki plate was placed on the modified adapter of *XtalCheck-S* plate reader attachment (Figure 1a) mounted on the goniometer omega stage. Two dozen of crystals were used for initial screening to rank diffraction quality. Omega and theta angles and then X, Y, and Z coordinates were adjusted in order to center crystals at the eucentric height of the X-ray focusing region. After centering, diffraction data was collected for each crystal (Figure 1b). Well-diffracting crystals were selected for further use in data collection and the exposure time was optimized to minimize radiation damage. The best diffracting crystals were grown in buffer containing 0.09 M HEPES-NaOH pH 7.5, 1.26 M sodium citrate tribasic dihydrate, 10% v/v glycerol (Crystal Screen Cryo (Cat#HR2-122)).

**Figure 1.**
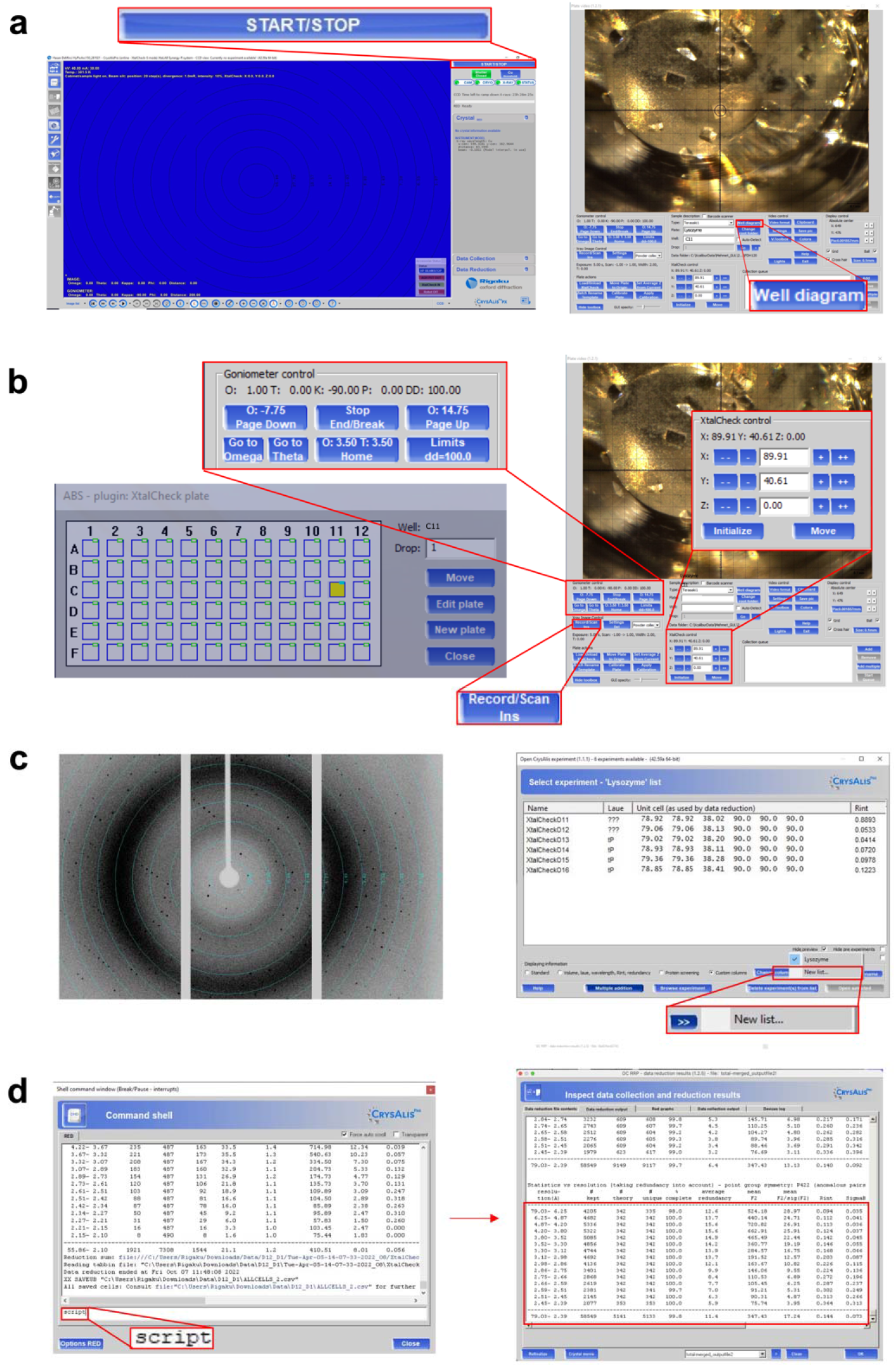
Workflow of structure determination with *XtalCheck-S* in *CrysAlisPro*. **(a)** After placing the crystallization plate to the goniometer and clicking the *START/STOP* button, camera starts to show the plate. **(b)** Out of 72, the desired well is selected through the well diagram button, and crystals are screened to collect diffraction data. Crystals are centered by changing the parameters from goniometer and XtalCheck control panels, and data collection is started by the *Record/Scan* button. **(c)** Diffraction data are obtained, and suitable data are added to the new list for data processing. **(d)** Data from each crystal are processed with the “`proffitbatch`” script. Then, the obtained data of different crystals are merged, and density map statistics are listed.

During data collection, *XtalCheck-S* was set to oscillate as much as the detector distance would allow in order to maximize crystal exposure oscillation angles. Diffraction data were collected (21 frames total) for 1 min and 45 sec (5 secs/frame) for each run from all individual crystals. A total of 13 crystals were used. The detector distance was set to 100.00 mm, the scan width to 1.00 degree oscillation and the exposure time to 5.00 sec per image (Supplementary Table 1).

### Data processing

Once plate screening parameters were optimized for all crystals, 21 degrees of data collection was performed for each prescreened/selected crystal (Supplementary Figure 1). All crystals were queued in CrysAlisPro for complete data collection. An optimal unit cell was chosen, and peak finding and masking were performed for the data collected (Supplementary Figure 2). A batch script was used generated with the `xx proffitbatch` command for cumulative data collection. The batch data reduction was run on CrysAlisPro Suite by the `script` command (Figure 1c). Data reduction produced a file that contains all integrated unmerged and unscaled data (*.rrpprof) for each dataset. For merging all datasets as a reflection data (*.mtz) file, the *proffit merge* process from the *Data Reduction* section on the main window of *CrysAlisPro* was used. Reduced datasets (*.rrpprof files) were then merged again using *proffit merge* as described. All data was refinalized, merged, and scaled with *aimless* and *pointless* implementation in *CCP4*^19,20^. Finally, the processed data were exported to *.mtz formats (Figure 1d) (please see <u>XtalCheck SOP</u>).

### Structure determination

The crystal structure of lysozyme was determined at ambient temperature in space group P4_3_2_1_2 by using the automated molecular replacement program *PHASER* ^21^ implemented in the *PHENIX* software package ^22^. A previously published X-ray structure was used as an initial search model (PDB ID: 3IJV ^23^). 3IJV structural coordinates were used for the initial rigid-body refinement within the *PHENIX*. After simulated-annealing refinement, individual coordinates and Translation/Libration/Screw (*TLS*) parameters were refined ^24,25^. Additionally, composite omit map refinement implemented in *PHENIX* was performed to identify potential positions of altered side chains, and water molecules. The final model was checked and rebuilt in *COOT* ^26^ while positions with a strong difference density were retained. Water molecules located outside of significant electron density were manually removed. All X-ray crystal structure figures were generated with *PyMOL* (Schrödinger, LLC) and *COOT*.

## RESULTS

### Ambient temperature lysozyme structure is determined at the Turkish Light Source

We determined chicken egg lysozyme structure to 2.39 Å resolution at ambient temperature using Rigaku’s XtaLAB Synergy Flow System XRD equipped with a modified *XtalCheck-S* Terasaki plate reader adaptor (Figure 2; Table 1). The lysozyme structure acquired from our diffraction data aligns well with our recently published cryogenic lysozyme structure (PDB: 7Y6A ^16^) with an RMSD value of 0.256 Å. The Ramachandran statistics for the allowed, favored, and outlier regions are 97.64%, 2.36%, and 0.00%, respectively. We obtained a well-defined electron density that reveals all aspects of the structure, including side chains and coordinated water molecules (Figure 2 & Supplementary Figure 3). The 129 amino acid structure consists of 8 alpha-helices and 2 beta-sheets (Figure 3 & Supplementary Figure 4).

**Figure 2.**
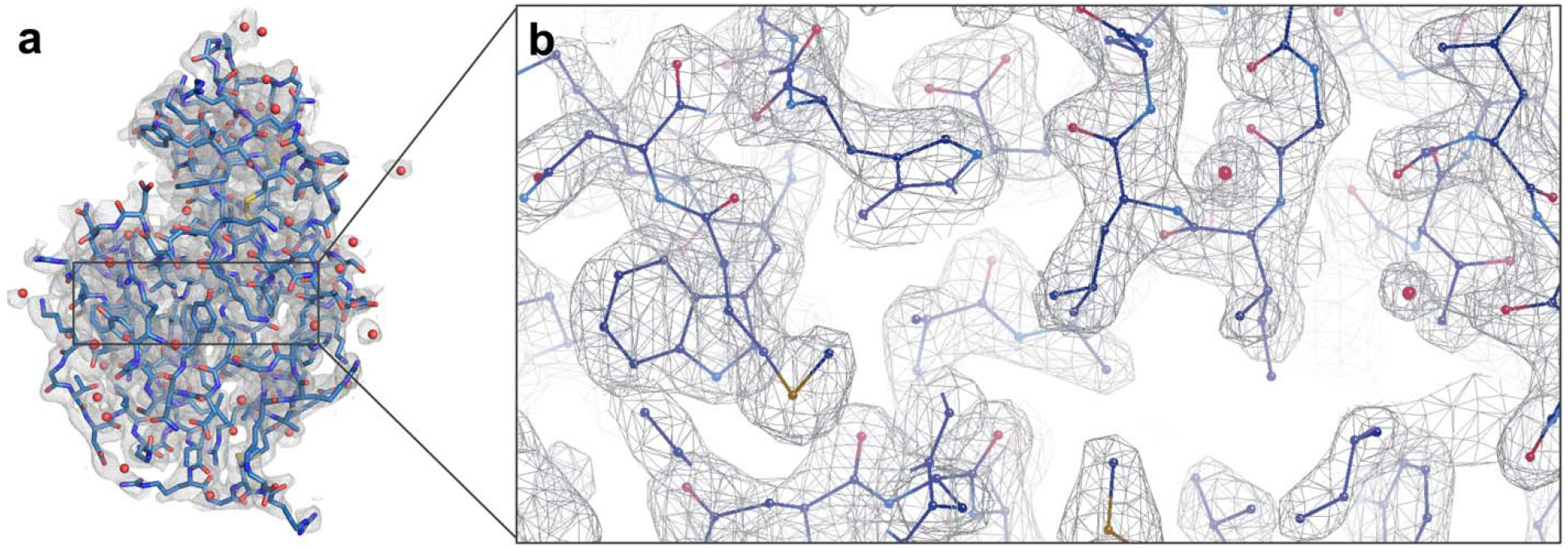
Ambient temperature structure of lysozyme (PDB ID: 8H3W). **(a)** 2Fo-Fc simulated annealing-omit map is shown in gray and contoured at 1.0 σ level. **(b)** The lysozyme structure is shown in the stick representation.

**Table 1.**
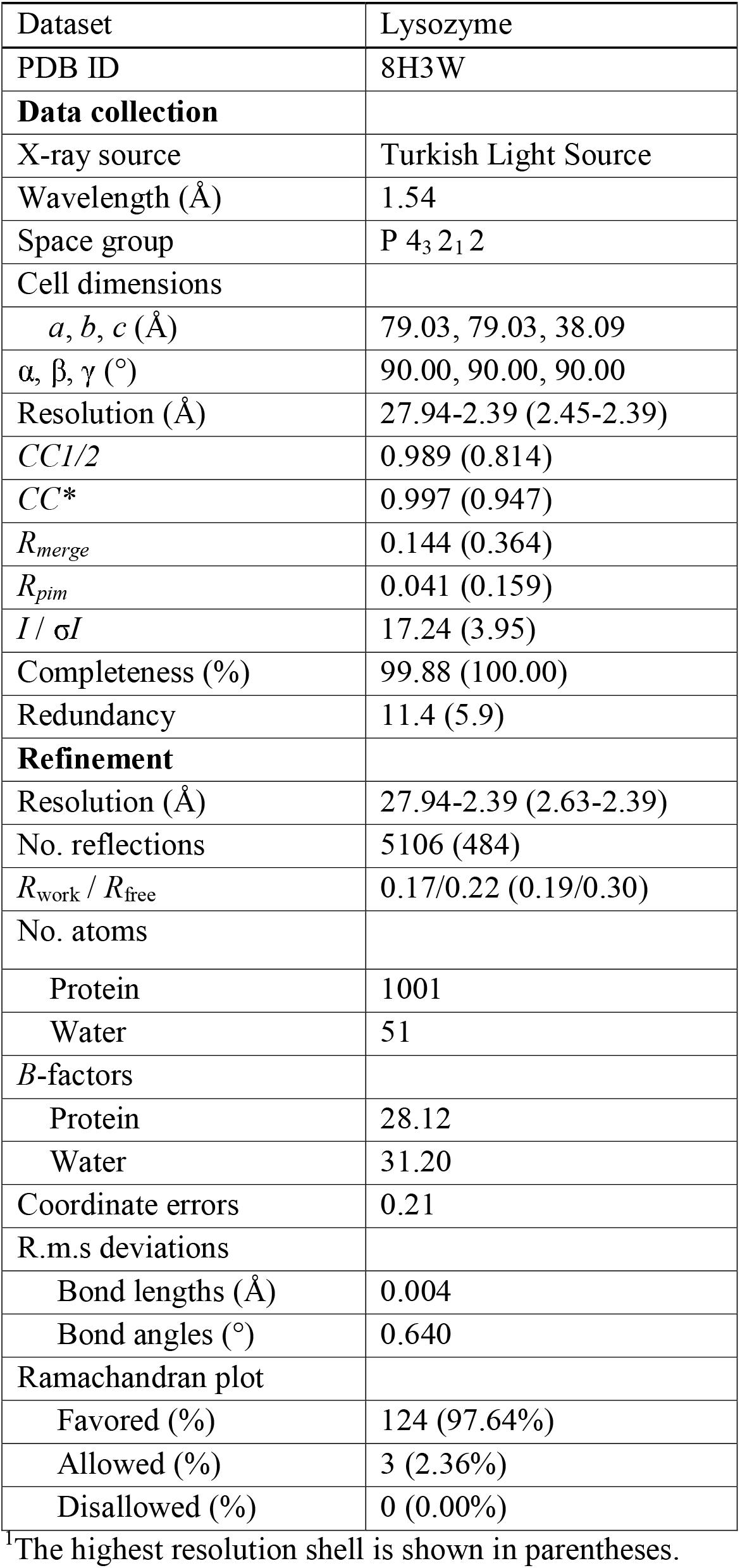
Data collection and refinement statistics.

**Figure 3.**
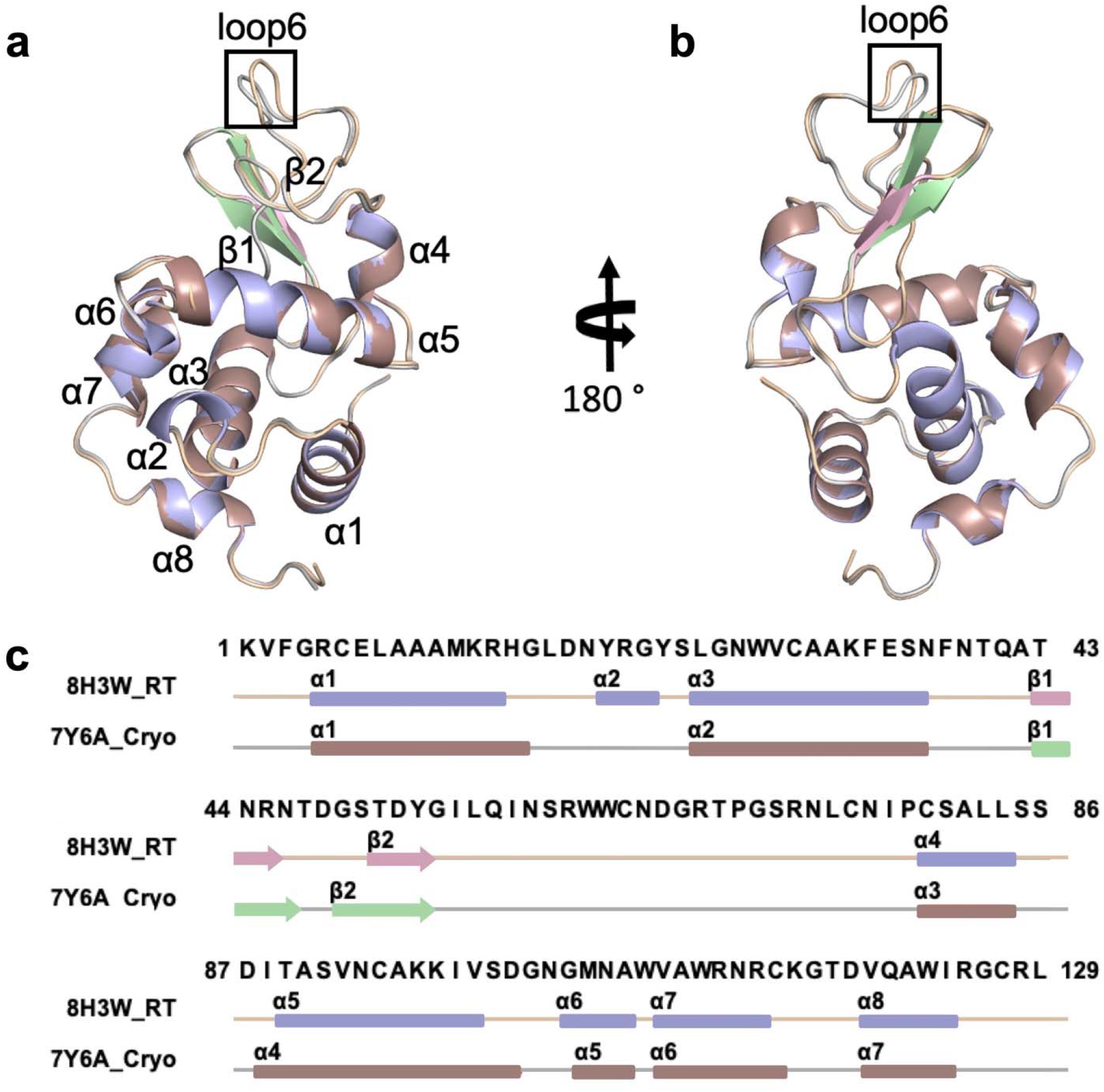
Secondary structure representation of chicken egg lysozymes. **(a-b)** The chicken egg lysozyme structure at ambient temperature (PDB ID: 8H3W) is superposed with the cryogenic structure (PDB ID: 7Y6A) with RMSD value of 0.256. Two side views are presented in the panel by rotating the structure 180 degrees on the y-axis. **(c)** Structure-based sequence alignment of lysozyme is indicated with secondary structures based on color code (alpha-helices: lightblue, darksalmon; beta-sheets: lightpink, palegreen; loops: wheat, gray, respectively).

### Lysozyme shows small structural changes at ambient temperature

The structure-based sequences of the cryogenic and ambient temperature lysozyme structures were aligned using *Jalview* ^27^ (Figure 3). Overall, the structures are almost identical with an additional mini-helix formation (□2) and slightly shorter beta-sheets in the ambient structure. A further comparison between cryogenic (PDB ID: 7Y6A ^17^) and ambient temperature lysozyme structures, we have has been made by examining B-factors (Supplementary Figure 5), suggesting that the side chains of the amino acids have greater mobility in the ambient temperature structure than in the cryogenic structure. Minor conformational changes were observed based on the comparison of secondary structures, with the exception of loop 6 (Supplementary Figure 6-11).

### Ambient temperature lysozyme displays lower radiation damage compared to cryogenic structure

Structural differences induced by radiation damage between cryogenic and ambient structures were compared using the *RABDAM* program ^28^. B_Damage_ and B_net_ values were calculated using the full atomic isotropic B-factor values of selected atoms and are presented in kernel density plots in Figure 4. The highest B_Damage_ value of 3.30 was observed on the Arg128 N atom (999) of the cryogenic lysozyme structure (PDB ID: 7Y6A ^17^) while in the ambient temperature structure (PDB ID: 8H3W), the highest B_Damage_ value (2.06) was observed on the Arg61 N atom (480) (Figure 4a). B_net_ values calculated for the Asp and Glu side chain oxygen atoms, for the 7Y6A structure is B_net_ =2.1 and median is 0.95, and for the 8H3W structure B_net_ = 2.6 and median is 0.97 (Figure 4b).

**Figure 4.**
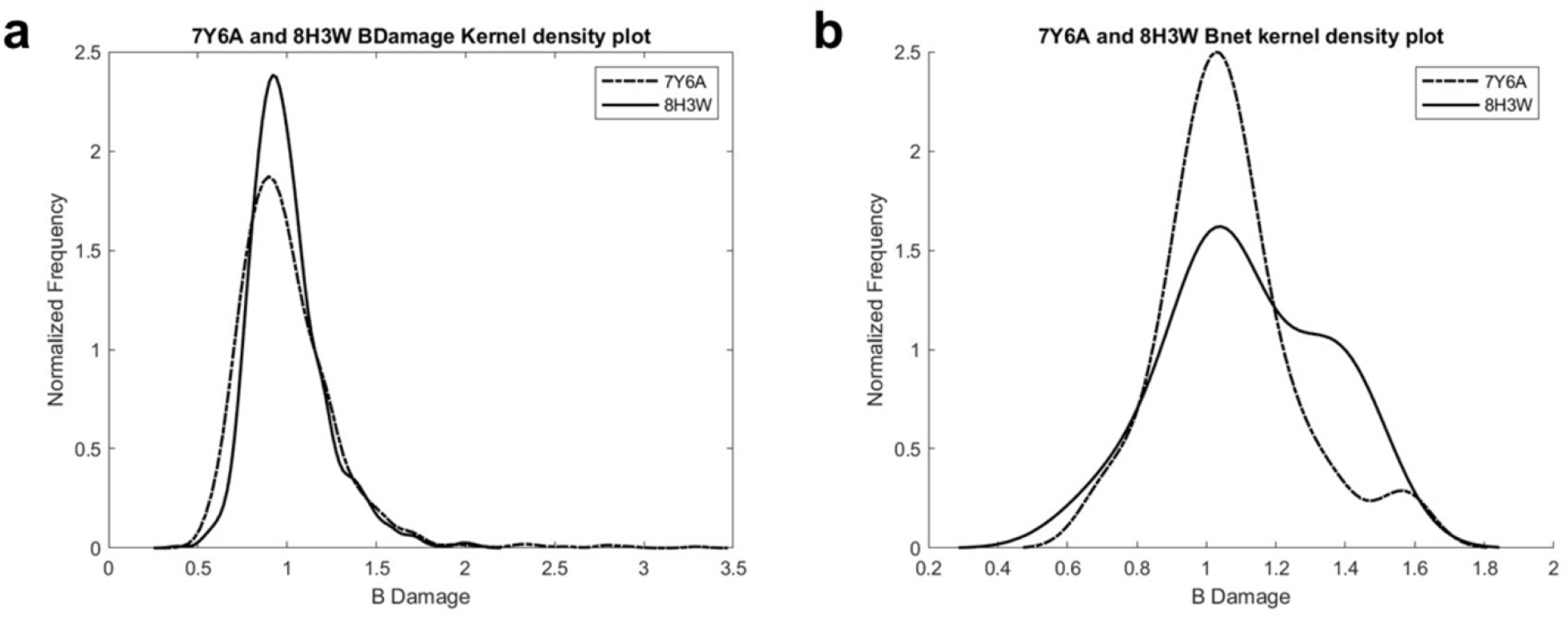
Values calculated using RABDAM software. **(a)** B_Damage_ distribution plots of the cryogenic lysozyme (PDB ID: 7Y6A) and the ambient temperature lysozyme (PDB ID: 8H3W)structures. **(b)** B_net_ distribution plots; B_net_ = 2.1 and median is 0.95 for 7Y6A structure, and **for** the 8H3W structure B_net_ = 2.6 and median is 0.97.

### XtaLAB Synergy Flow system: XtalCheck-S provides a user-friendly platform for ambient data collection

*Turkish DeLight* switched to *Warm* data collection mode is equipped with a Hybrid Photon Counting X-ray detector (HyPix-Arc 150°), high-performance X-ray source and a goniometer-mountable plate reader module that can be remotely controlled by *CrysAlisPro* software. Our modified *XtalCheck-S* platform employing affordable Terasaki plates is a low-cost, user-friendly, and automated *in situ* alternative crystallography technique that enables the screening, collection and data processing from *multiple* protein crystals in a single crystal X-ray diffractometer (SC-XRD) home-source from a series of wells on a single plate (Figure 5a). Modified *XtalCheck-S* is a highly versatile tool for *in situ* screening and data collection from protein crystals, small molecules, and powder samples. We have adapted this module for 72-well Terasaki plates for use in place of the specially designed 96-well plate unique to *XtalCheck-S* for both macromolecule and small molecule data collection.

**Figure 5.**
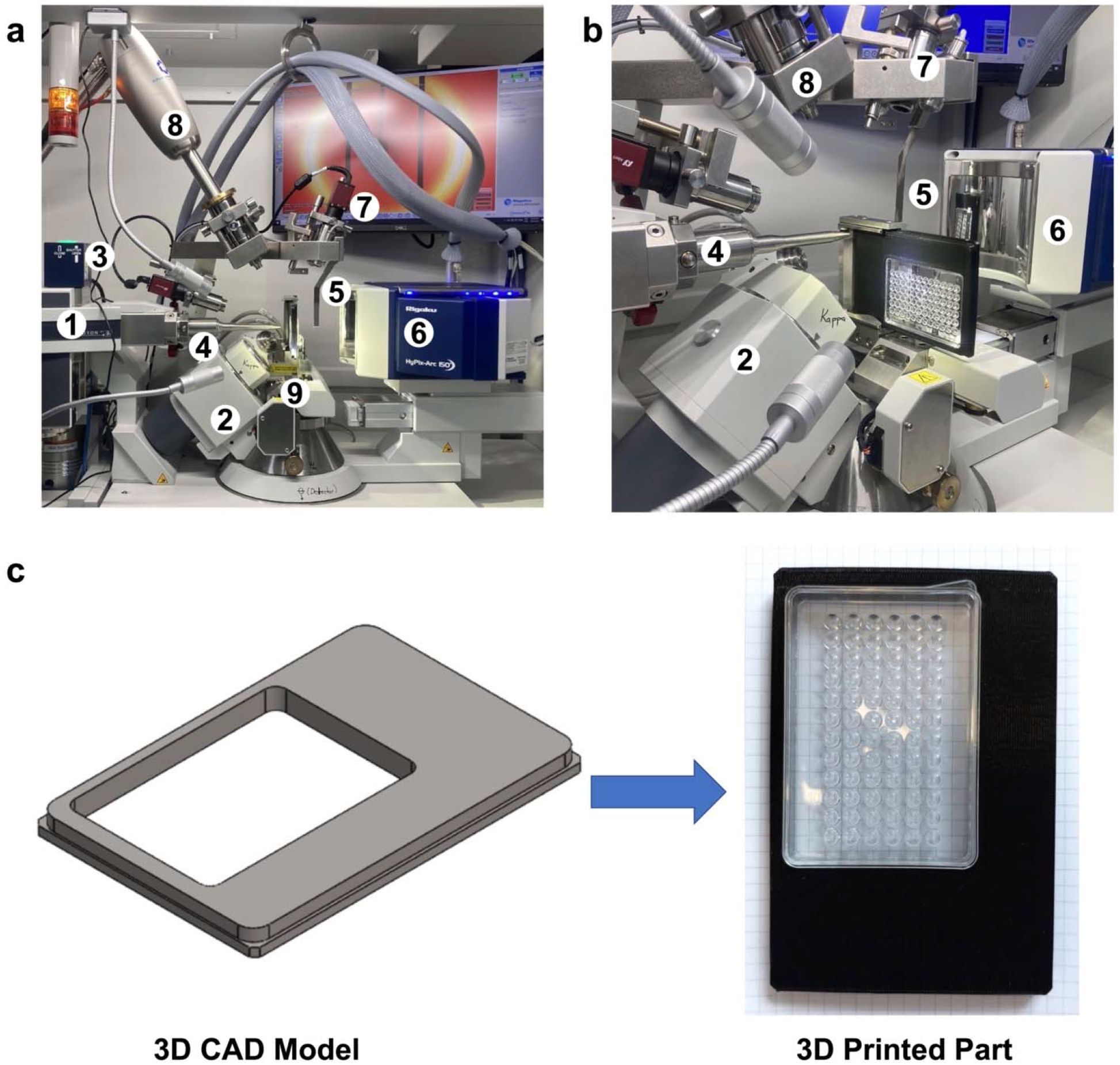
Overview of XtaLAB Synergy Flow system. **(a, b)** XtalCheck-S module. **(1)** X-ray source; **(2)** four-circle Kappa goniometer; **(3)** shutter; **(4)** collimator; **(5)** beamstop; **(6)** X-ray detector; **(7)** video microscope; **(8)** low temperature attachment; **(9)** XtalCheck module. **(c)** 3D modeled plate holder adapter and 3D printed part.

We designed a Terasaki plate holder adapter and printed it with a 3D printer (Replicator+, Makerbot, NY) that encloses the Terasaki plate (Figure 5c). A 3D printable .stl file of the plate holder is available in the Supplementary Files (Supplementary Material Plate_Holder.stl file). Previously added paraffin oil, a viscous material, significantly slows down the sliding of the protein crystals off the vertically mounted Terasaki plate. After gently placing the Terasaki plate in the plate holder, it is carefully slid into the plate holder mount on the goniometer stage (Figure 5b). Necessary parameters can be manipulated by the *XtalCheck-S* system by accessing the *plate video* panel over the *CrysAlisPro* software. From here, the plate is labeled under description as ‘Lysozyme’, our custom-made ‘Terasaki plate’ is selected as the plate type, and the *well diagram* button is pressed in order to select a well with crystals to be screened (Figure 1a, b). The crystal focus is provided by centering the crystal using the *goniometer control* and *XtalCheck-S control* points in the *plate video* panel (Figure 1b). Protein crystals are first checked with the *powder diffraction* option on the *X-ray image control* by editing the *powder collection parameters* panel (i.e., theta, omega degrees and exposure time). The *powder diffraction* option is more convenient for both protein crystallography (PX) and chemical crystallography (CX) than other options due to simpler and faster screening (Figure 1b). The *Settings/Del* button allows one to modify crystal “screening” parameters, while the *Record/Scan* button sets up data collection for crystal in view (Figure 1b). As long as all crystals have similar unit cells, hundreds of data sets can be collected from tens of wells in a single plate with the *multiwell-multicrystal* approach. Cumulative data reduction can be performed through the easy-to-use GUI or by using a simple script to generate a merged *.mtz file (see XtalCheck SOP).

## DISCUSSION

X-ray crystallography, cryo-electron microscopy (Cryo-EM), mass spectrometry (MS), nuclear magnetic resonance spectroscopy (NMR), electron paramagnetic resonance spectroscopy (EPR), and small angle X-ray scattering (SAXS) are established techniques for the investigation of the structure and dynamics of biomacromolecules. X-ray crystallography is the most popular and robust among these techniques for structure determination owing to short X-ray wavelengths and diffraction properties suitable for high-resolution protein structure determination ^28^. X-ray crystallography can provide insights on macromolecular dynamics at ambient temperature, especially when combined with serial data collection ^30^. Although this is the case, sample size and volume in addition to managing structural radiation damage when using these techniques is a challenge. Cryogenic single-crystal XRD approaches mitigate radiation damage; however, they can provide only limited protein dynamics information ^30,31^. Additionally, cryoprotectants such as glycerol, MPD, ethylene glycol, and PEG may also result in significant increase in crystal mosaicity. Moreover, flash-freezing during cryogenic sample preparation can cause the contraction of protein crystals due to lattice repacking and the disruption of intra- and intermolecular contact interfaces ^32^. Ambient temperature SFX performed at XFELs and serial millisecond crystallography (SMX) performed at synchrotrons provide new strategies for addressing these issues. In particular, fourth generation XFELs provide extremely short X-ray pulses and are a billion times brighter than any other current X-ray sources, facilitating completely different approaches to structure determination ^30,33^. Radiation damage on small-sized crystals can be prevented with the aid of a continuous sample delivery system that supplies a fresh crystal for each pulse, which is known as the “diffract-and-destroy” concept in SFX ^34^. Thus diffraction data are obtained from nano- or micro-sized crystals that are streamed across the X-ray beams using a fixed-target or a liquid jet system ^30,31,33^. However, serial crystallography (SX) techniques can be more challenging than conventional X-ray crystallography, due to a considerable number of crystals being consumed once crystal samples are exposed to X-rays ^31,33^. Hence, numerous research groups prefer to use their primary home-source XRD to screen their crystals or collect data. Therefore, there is a significant demand for easy-to-use and efficient XRD infrastructures where optimum crystal data collection and processing procedures can be realized ^17^.

*XtalCheck-S* is a user-friendly goniometer-mountable attachment for serial scanning and “serial” ambient temperature data collection of various types of samples including protein crystals, small molecule crystals and powder samples in a 72-well Terasaki plate (Figure 5). Protein crystallography often requires screening large numbers of crystals to identify the best diffracting crystal. This module can differentiate between a salt and a protein crystal in seconds without necessitating freezing the crystals. It is fully automated and suited to collect diffraction data directly from a Terasaki plate with reduced background noise. Every step from the centering of the crystals to the collection of the diffraction data is easily traceable, measurable, and viewable remotely (Figure 1). A large quantity of datasets at ambient temperature can be collected from a single plate and multiple wells in minutes. Serially collected data from thousands of crystals is combined with the single crystal data principle. Thus, it provides complete data sets that can be used for structure determination obtained through this module, offering a distinct solution to SX.

In this study, we collected lysozyme diffraction data for up to 1.5 minutes for each run and 20 minutes total, using the *XtalCheck-S* module and determined the lysozyme structure at 2.39 □ resolution with 100% completeness (Figure 2). We have confirmed that the ambient-lysozyme structure closely matches with the cryogenic-lysozyme structure that we have published recently (Figure 3) (RMSD: 0.256 A, PDB ID: 7Y6A ^17^). Compared to the 7Y6A ^17^ lysozyme structure, we observed minor conformational changes and more flexibility with increased B-factors, suggesting slightly more plasticity than the cryogenic lysozyme structure, as expected (Figure 3; Supplementary Figure 5; Supplementary Figure 6; Supplementary Figure 12). Moreover, the radiation damage differences between the cryogenic (7Y6A ^17^) and ambient (8H3W) temperature structures determined using the same home-source XRD (*Turkish DeLight*) indicate that the overall B_Damage_ (all atom calculation of B_Damage_ values) value of the ambient structure (2.06) was less than our cryogenic structure (3.30), suggesting less radiation damage occurred (Figure 4a).

Collectively, we have presented the beyond-the-state-of-the-art *XtalCheck-S* module configured with a user-friendly *CrysAlispro* software suite in *Turkish DeLight*. The diffraction data of the *in situ* lysozyme structure determined in this study was cost-effectively collected in a noticeably short time at ambient temperature with the single plate *multiwell-multicrystal* principle and reduced radiation damage when compared to the data collection for the cryogenic structure. Accordingly, *Turkish DeLight* offers a novel perspective on traditional SX, allowing rapid, robust, and simple micro-batch data collection from multiple crystals over multiple wells.

## Data availability

The lysozyme structure in this article has been deposited to the Protein Data Bank under the accession number 8H3W. Any remaining information can be obtained from the corresponding author upon reasonable request.

## Acknowledgement/Disclaimers/Conflict of interest

Authors would like to dedicate this manuscript to the memory of Dr. Albert E. Dahlberg and Dr. Nizar Turker. The authors gratefully acknowledge use of the services and facilities of the Koç University Isbank Infectious Disease Center (KUISCID). H.D. acknowledges support from NSF Science and Technology Center grant NSF-1231306 (Biology with X-ray Lasers, BioXFEL). A.K. acknowledges support from Scientific and Technological Research Council of Türkiye (TÜBİTAK, 2218 - National Postdoctoral Research Fellowship Program under project number 118C476). B.V.K. are funded by TÜBİTAK 2232 International Outstanding Researchers Program (Project No: 118C225). This publication has been produced benefiting from the 2232 International Fellowship for Outstanding Researchers Program, 2236 CoCirculation2 program and the 1001 Scientific and Technological Research Projects Funding Program of the TÜBİTAK (Project Nos. 118C270, 121C063 and 120Z520). However, the entire responsibility of the publication belongs to the authors of the publication. The financial support received from TÜBİTAK does not mean that the content of the publication is approved in a scientific sense by TÜBİTAK. Coordinates of the lysozyme structure have been deposited in the Protein Data Bank under accession code 8H3W.

## Author contributions

H.D. designed the experiments. M.G., E.A., E.D., J.A.J., A.S., A.Kepceoglu, M.Y., F.B.E., I.Y., B.T., N.B., N.T., Z.N., G.K., S.S.P., C.K., C.K.F., O.G., N.A., E.K.A., H.Cetinok, N.E.A., K.S., B.A., S.Tavli, H.G., H.Ciftci, B.K.T., B.V.K., and H.D. performed sample preparation and crystallization. M.G., E.A., E.D., J.A.J., A.S., A.Kepceoglu, M.Y., F.B.E., I.Y., B.T., N.B., N.T., Z.N., G.K., S.S.P., C.K., C.K.F., O.G., N.A., E.K.A., H.Cetinok, N.E.A., K.S., B.A., S.Tavli, H.G.,

S.A., H.Ciftci, B.K.T., B.V.K., A.Kati, and H.D. executed data collection. M.G., E.A., E.D., J.A.J., A.S., A.Kepceoglu, M.Y., F.B.E., I.Y., B.T., N.B., N.T., Z.N., G.K., S.S.P., C.K., C.K.F., O.G., N.A., E.K.A., H.Cetinok, N.E.A., K.S., B.A., S.Tavli, H.G., H.Ciftci, B.K.T., B.V.K., and H.D. performed data processing and structure refinement. The manuscript was written and prepared by M.G., E.A., E.D., J.A.J., A.S., A.Kepceoglu, M.Y., F.B.E., I.Y., B.T., N.B., N.T., Z.N., G.K., S.S.P., C.K., C.K.F., O.G., N.A., E.K.A., H.Cetinok, N.E.A., K.S., B.A., S.Tavli, H.G., S.A., M.O., M.F., S.Tekin, H.Ciftci, S.D., E.K., B.K.T., B.V.K., A.Kati, and H.D.

