## Supplementary Table and Figures for "Rapid and Efficient Ambient Temperature X-ray Crystal Structure Determination at Turkish Light Source"

**Supplementary Table-1**: Three different run parameters.

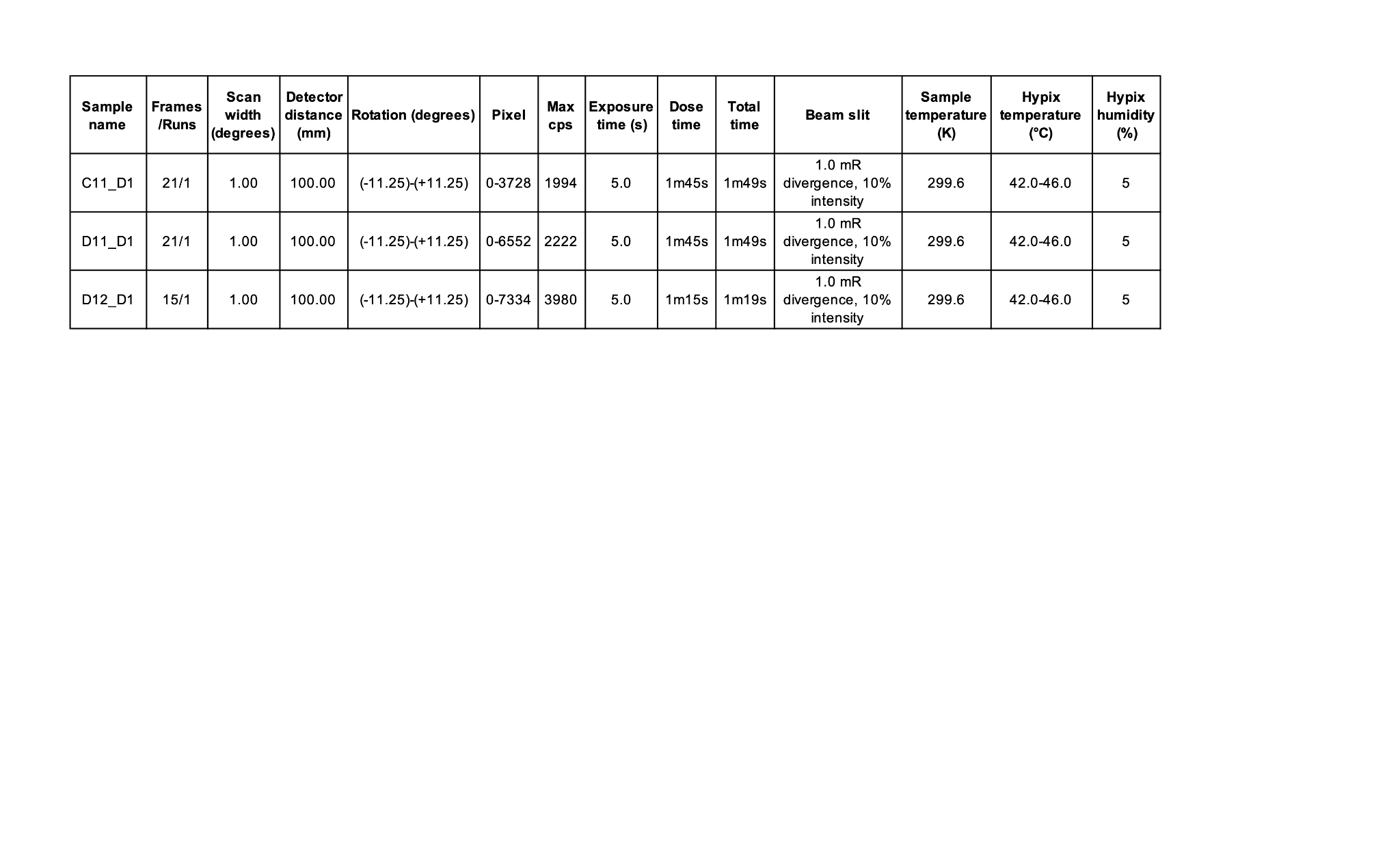

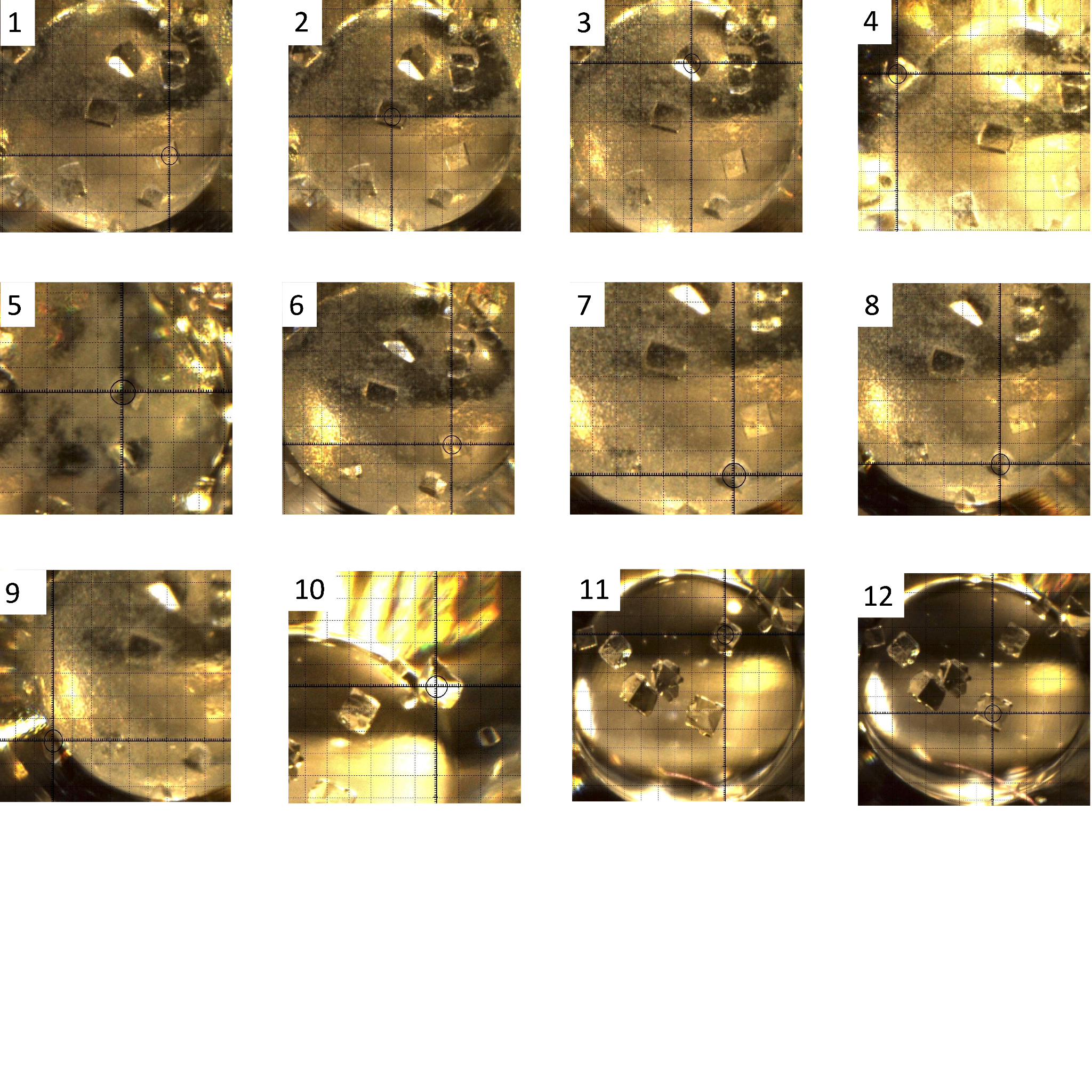

**Supplementary Figure 1.** Multiple crystals for data collection experiments.

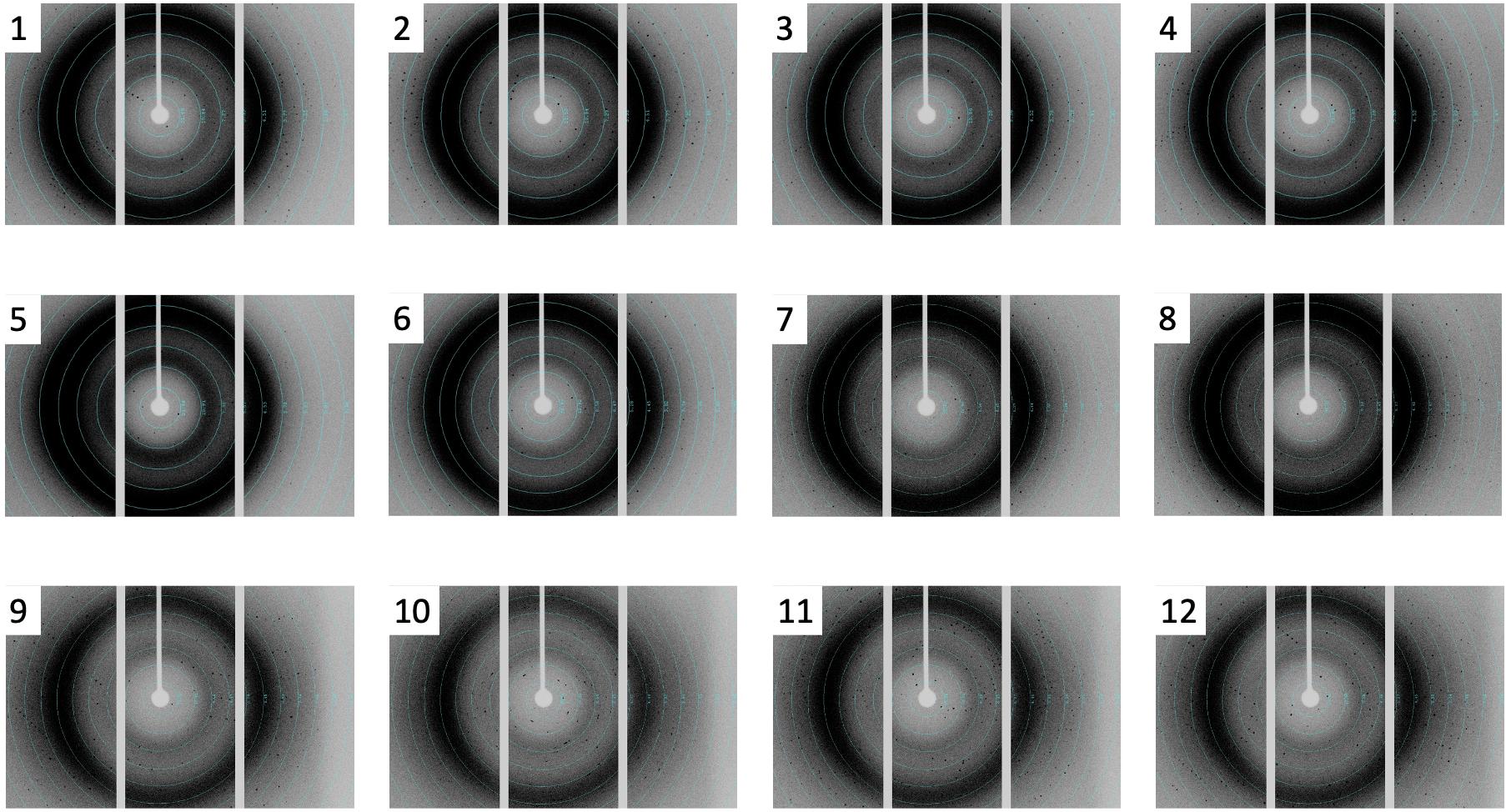

**Supplementary Figure 2.** Diffraction patterns collected from each crystal.

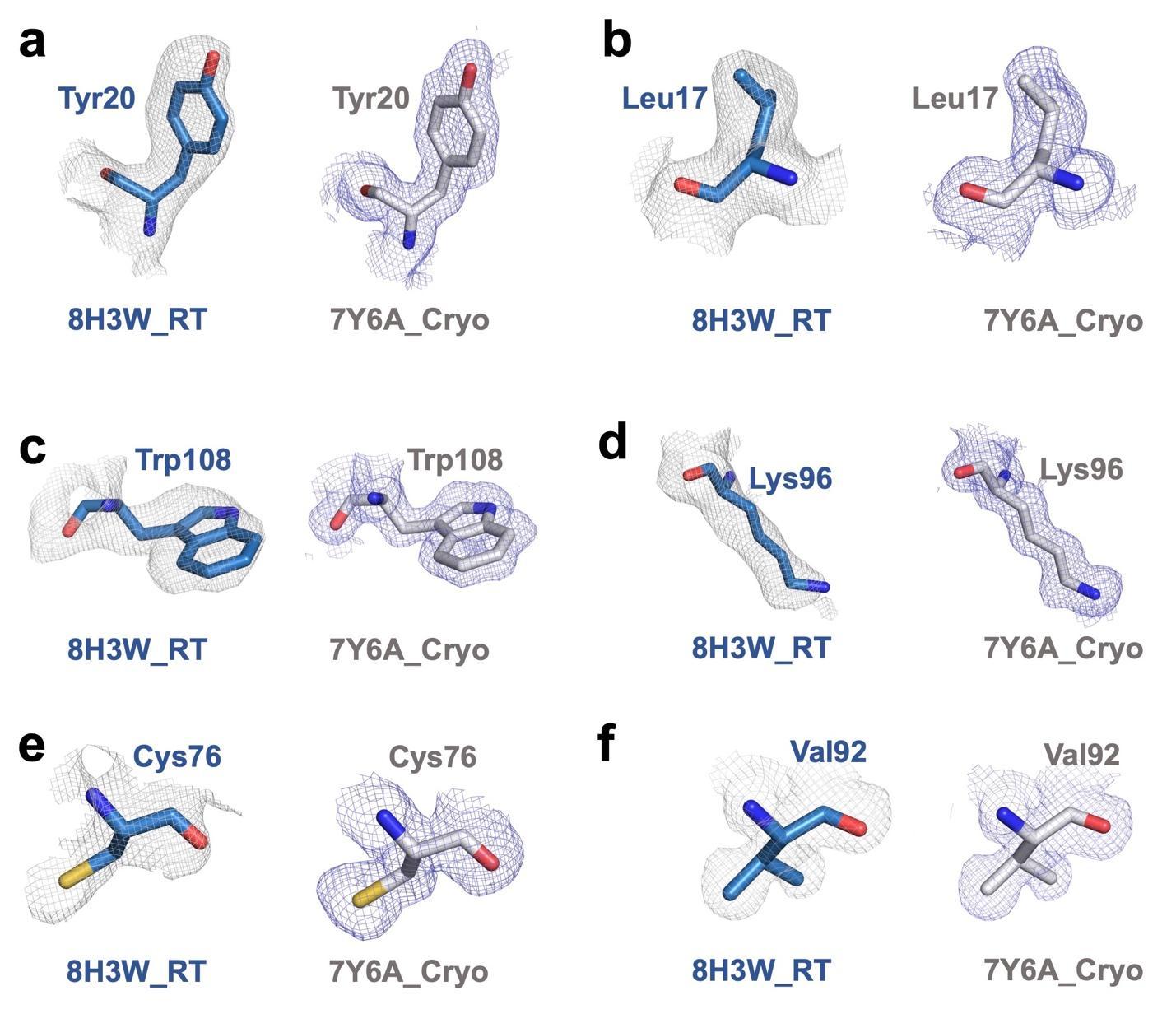

**Supplementary Figure 3.** Comparison of electron density map of ambient temperature (PDB ID: 8H3W) and cryogenic temperature (PDB ID: 7Y6A) chicken egg lysozyme structure for residues. 2Fo-Fc simulated annealing-omit map for ambient temperature structure is shown in gray while 2Fo-Fc simulated annealing-omit map for cryogenic structure is shown in slate.

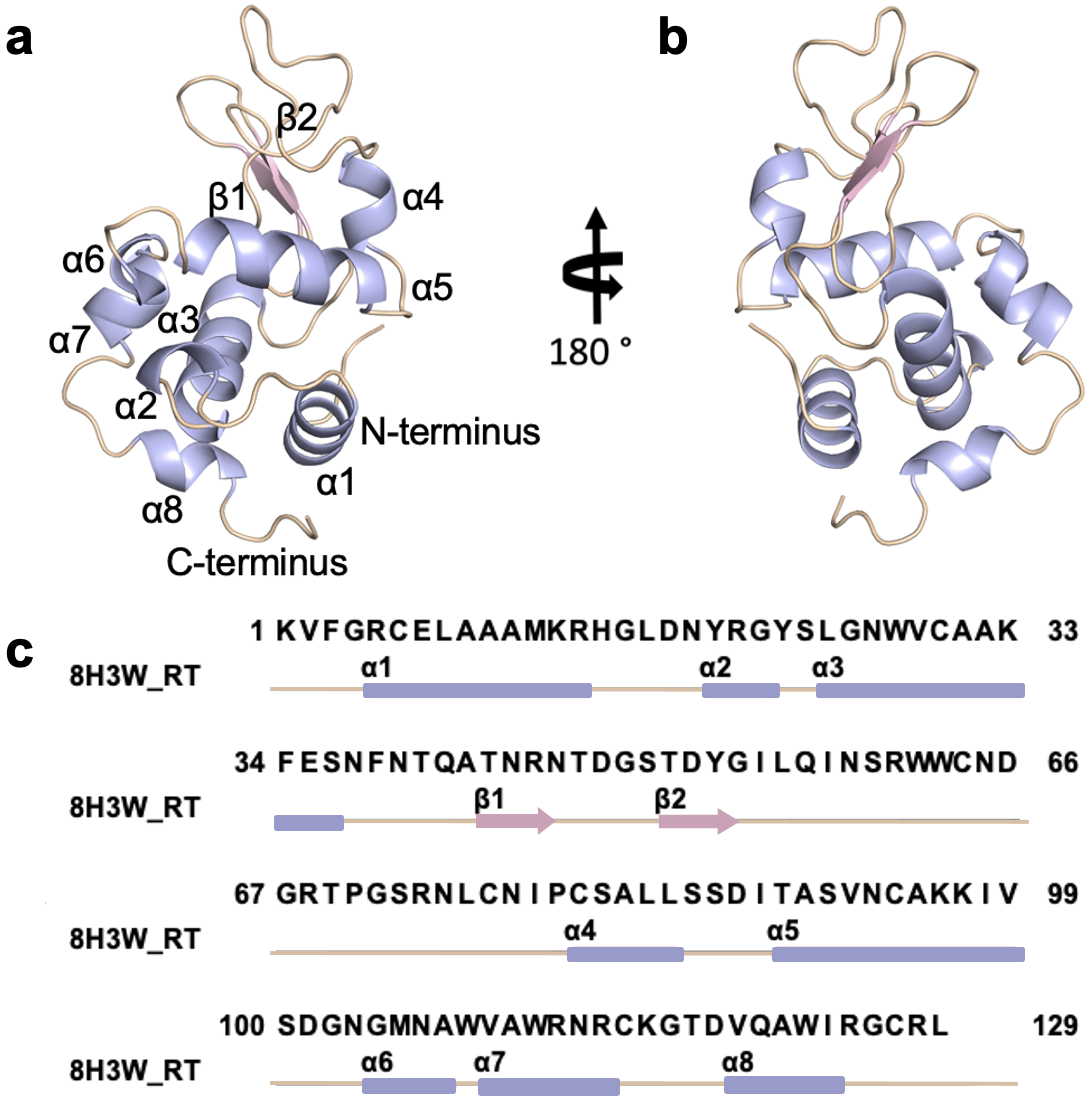

**Supplementary Figure 4.** Secondary structure representation of chicken egg lysozyme at ambient temperature (PDB ID: 8H3W). **(a-b)** The chicken egg lysozyme structure at ambient temperature is shown with cartoon representation. Two side views are presented in the panel by rotating the structure 180 degrees on the y-axis. **(c)** Structure-based sequence alignment of lysozyme is indicated with secondary structures based on color code (alpha-helices: lightblue; beta-sheets: lightpink; loops: wheat).

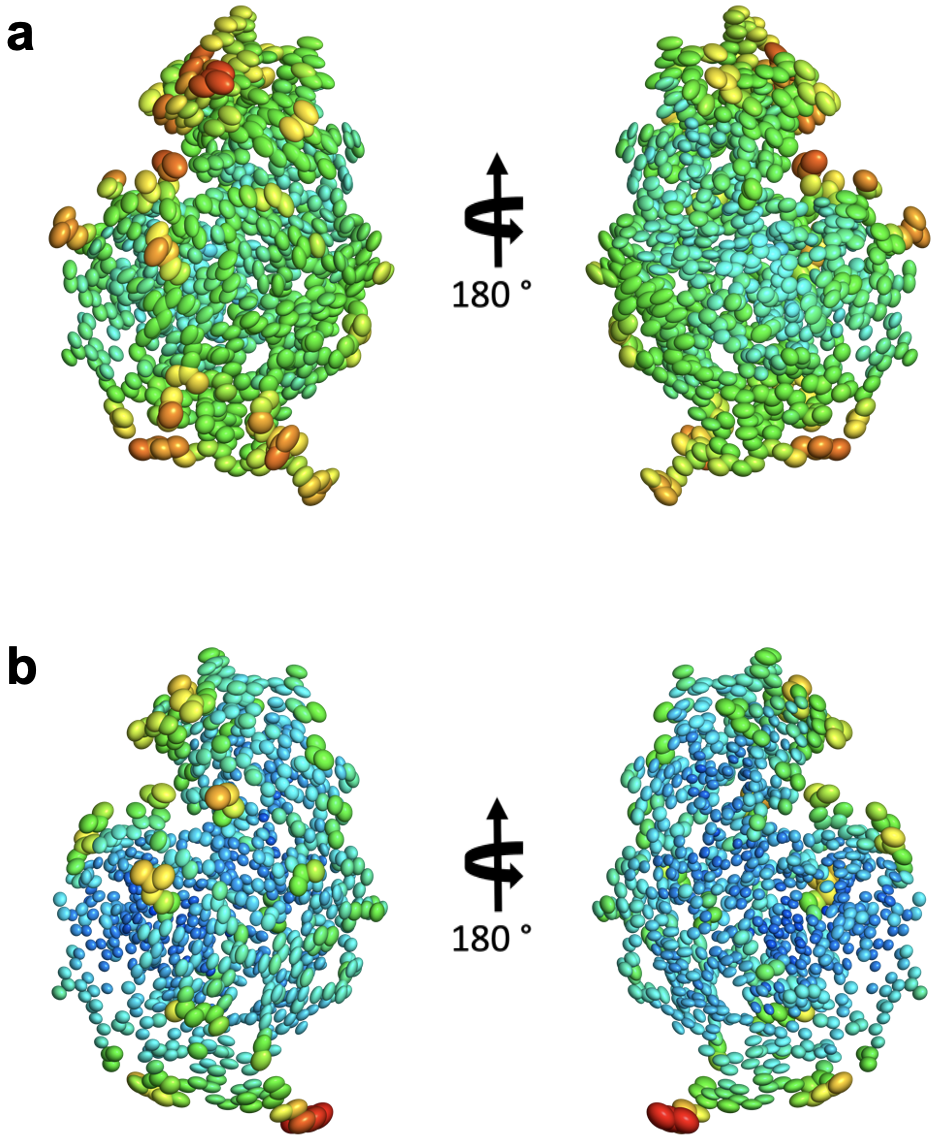

**Supplementary Figure 5.** Ellipsoid presentation of chicken egg lysozyme at **(a)** ambient and **(b)** cryogenic temperature.

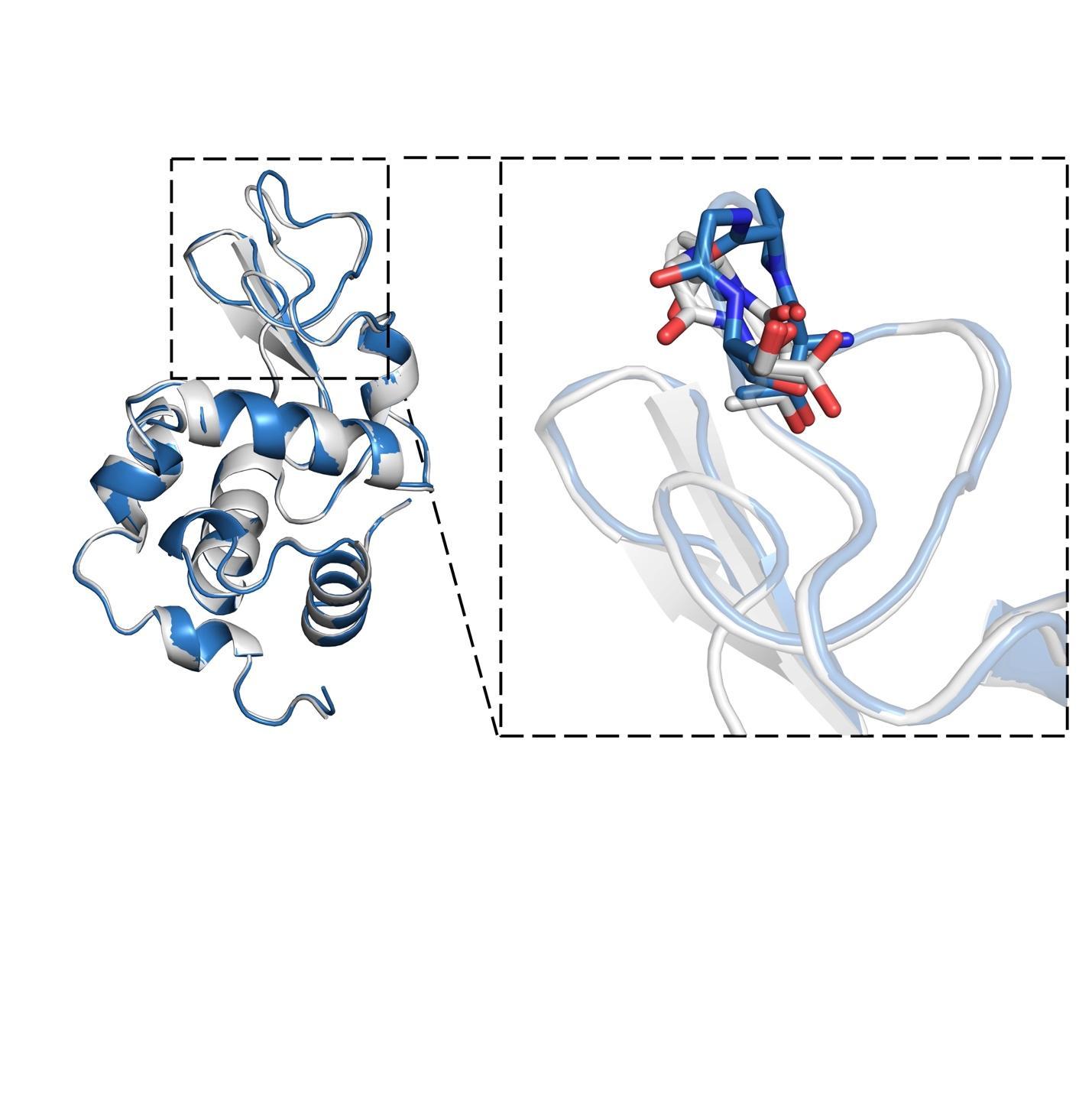

**Supplementary Figure 6.** Conformational changes on loop6. Ambient temperature lysozyme is shown in skyblue. Cryogenic lysozyme is shown in gray.

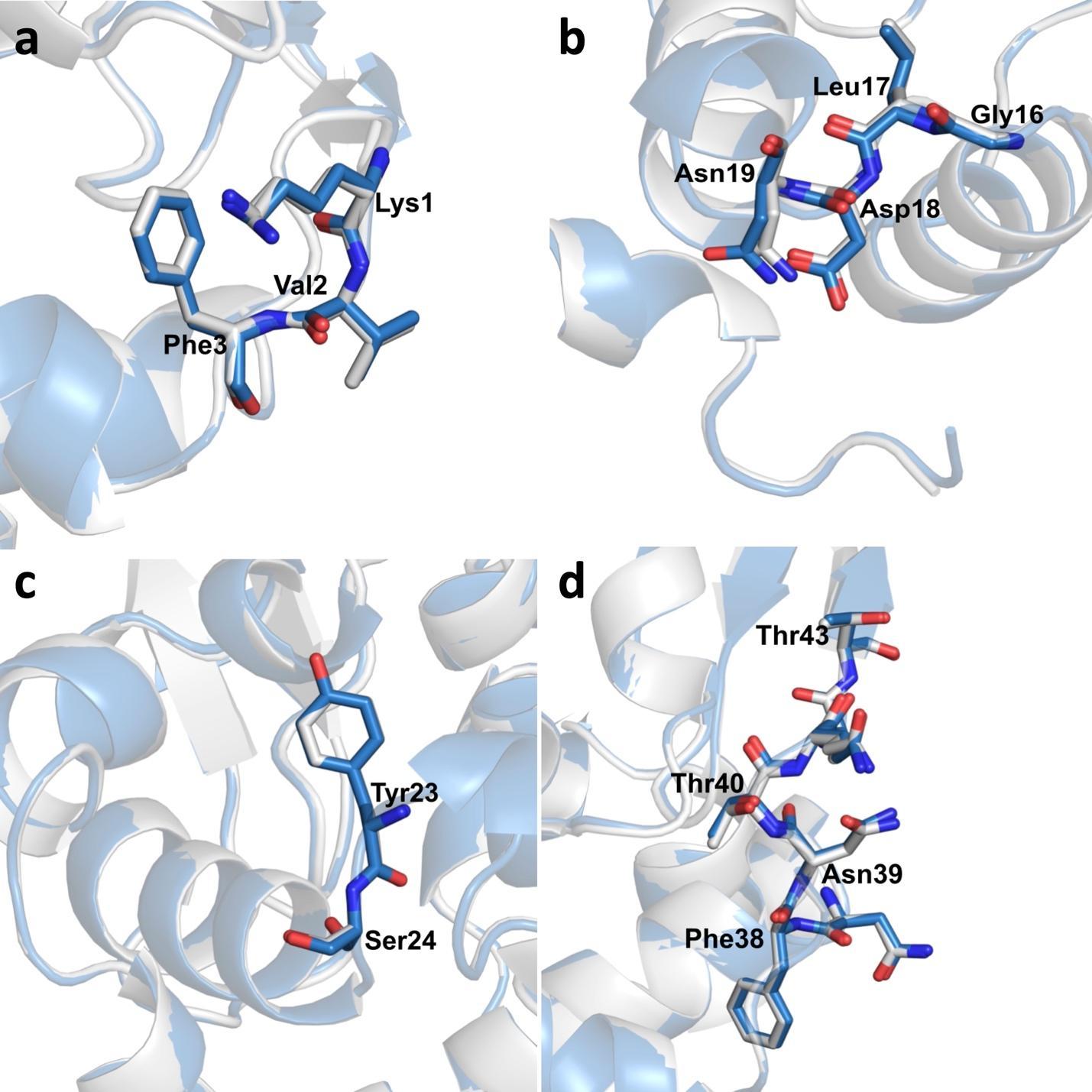

**Supplementary Figure 7.** Loop region (1-4) comparisons of ambient temperature lysozyme (skyblue) with cryogenic lysozyme (gray). RMS values are shown in parentheses. **(a)** Loop 1 (*0.109 Å*); **(b)** Loop 2 (*0.117 Å***)**; **(c)** Loop 3 (*0.001 Å*); **(d)** Loop 4 (*0.091 Å*).

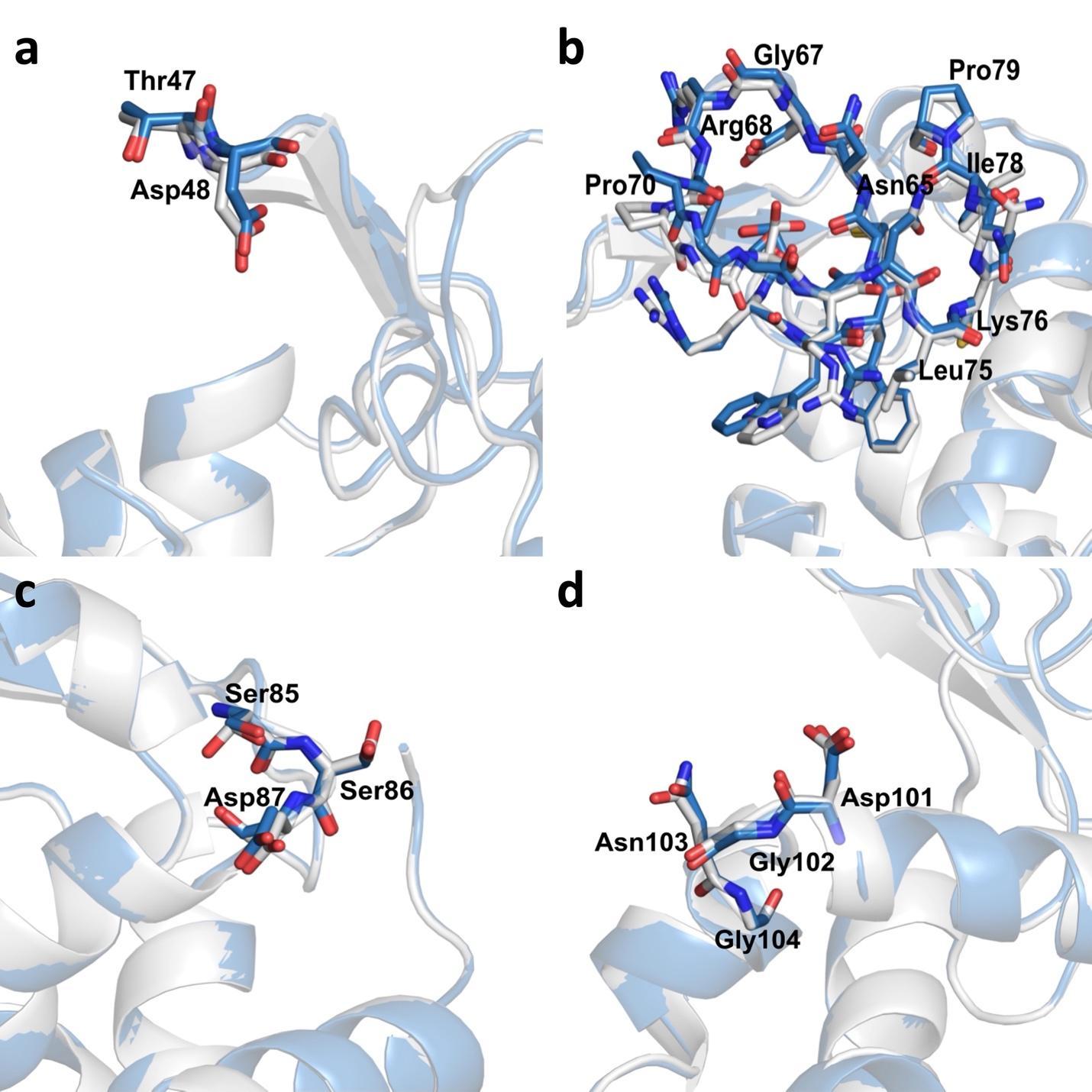

**Supplementary Figure 8.** Loop region (5-8) comparisons of ambient temperature lysozyme (skyblue) with cryogenic lysozyme (gray). RMS values are shown in parentheses. **(a)** Loop 5 (*0.235 Å*); **(b)** Loop 6 (*0.066 Å*); **(c)** Loop 7 (*0.066 Å*); **(d)** Loop 8 (*0.095 Å*).

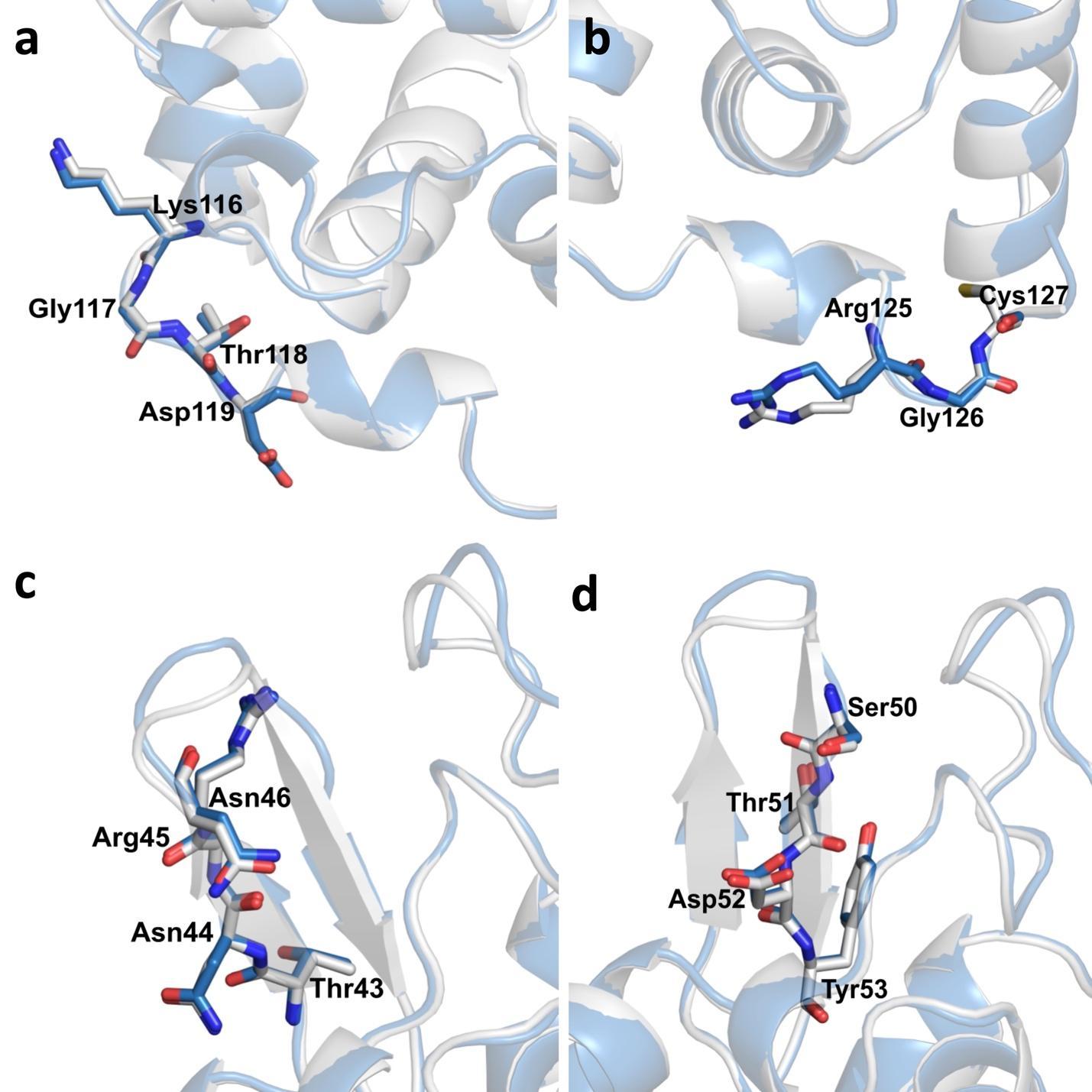

**Supplementary Figure 9.** Loop and beta-sheet region comparisons of ambient temperature lysozyme (skyblue) with cryogenic lysozyme (gray). RMS values are shown in parentheses. **(a)** Loop 9 (*0.052 Å*); **(b)** Loop 10 *(0.045 Å*) ; **(c)** Beta sheet 1 (*0.022 Å*) ; **(d)** Beta- sheet 2 (*0.066 Å*).

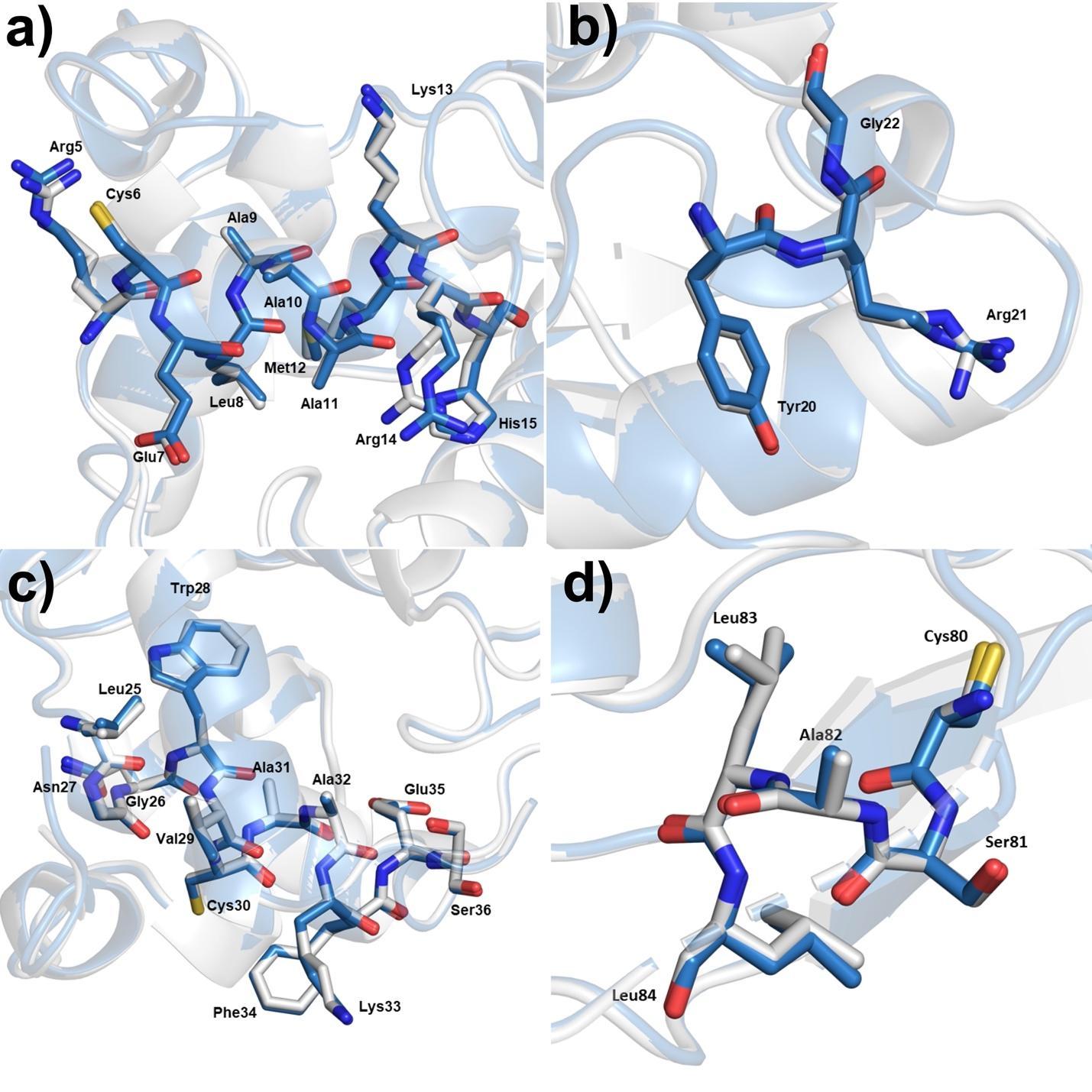

**Supplementary Figure 10.** Helix region (1-4) comparisons of ambient temperature lysozyme (skyblue) with cryogenic lysozyme (gray). RMS values are shown in parentheses. **(a)** Helix 1 (*0.097 Å*); **(b)** Helix 2 (*0.092 Å*); **(c)** Helix 3 (*0.070 Å*); **(d)** Helix 4 (*0.061 Å*).

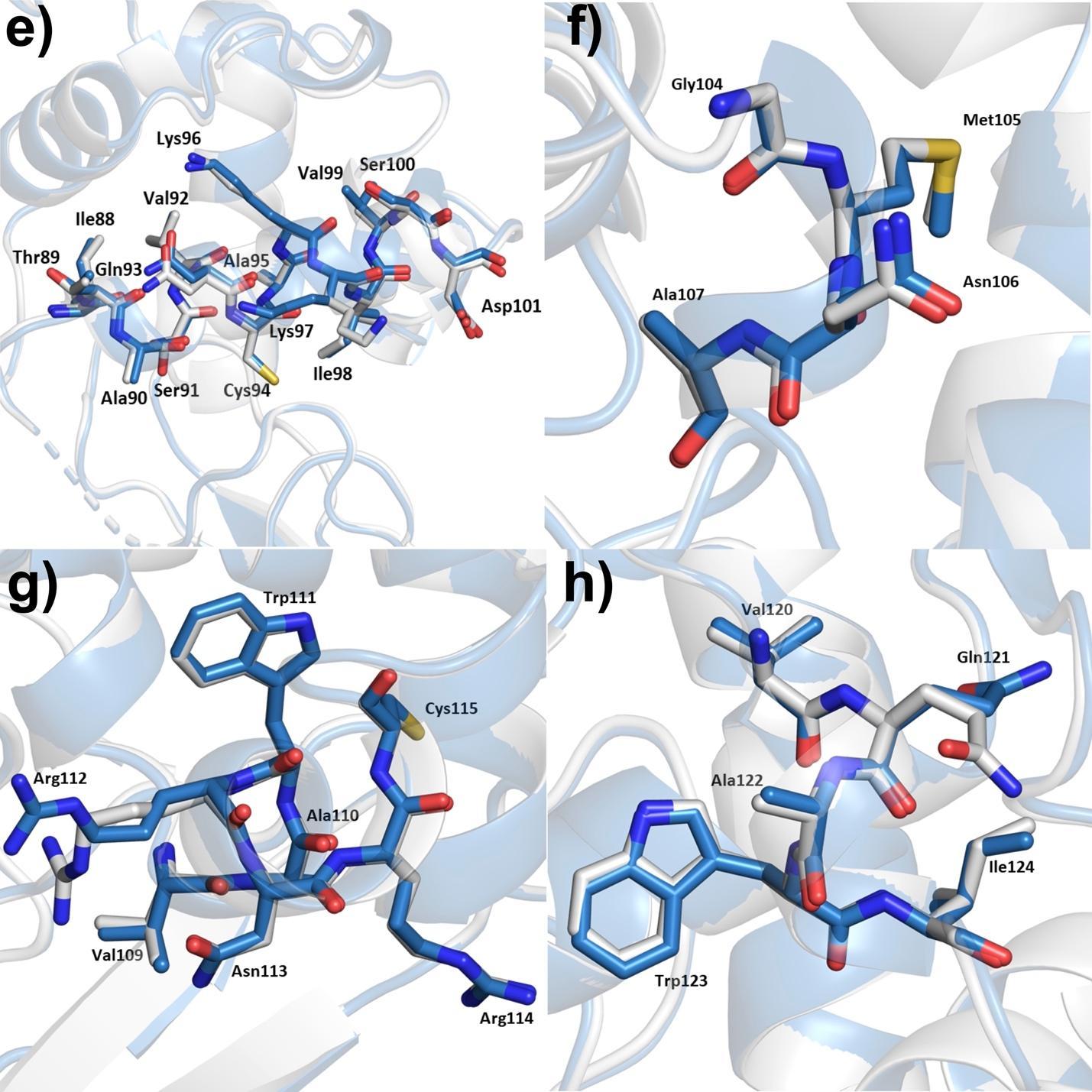

**Supplementary Figure 11.** Helix region (5-8) comparisons of ambient temperature lysozyme (skyblue) with cryogenic lysozyme (gray). RMS values are shown in parentheses. **(a)** Helix 5 (*0.100 Å*); **(b)** Helix 6 (*0.101 Å*); **(c)** Helix 7 (*0.115 Å*); **(d)** Helix 8 (*0.087 Å*).

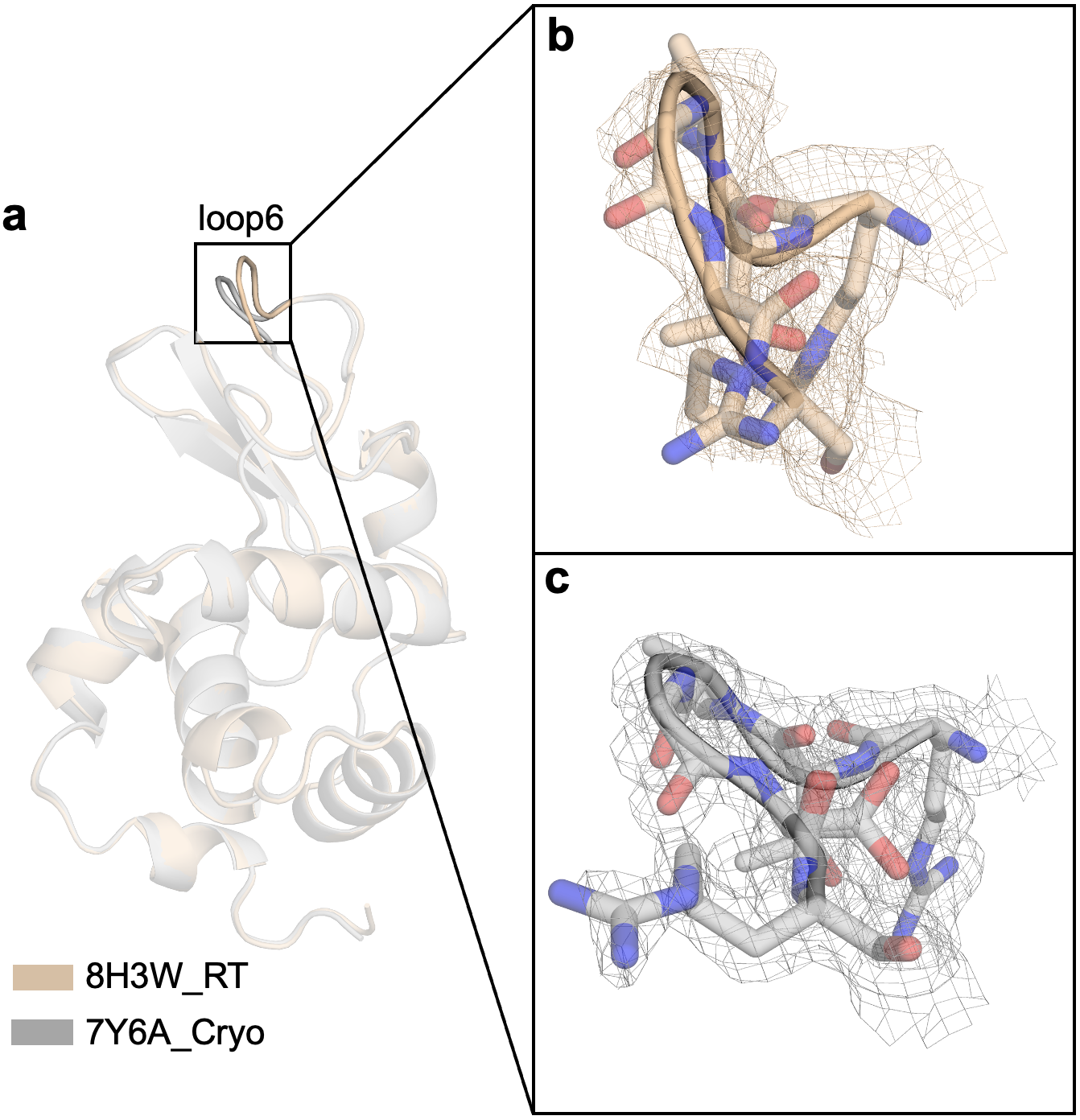

**Supplementary Figure 12.** The superposition of room and cryogenic temperature structure of lysozyme. **(a)** The room temperature structure of lysozyme is superposed with the cryogenic temperature structure of lysozyme with an RMSD value of 0.256. **(b)** 2Fo-Fc simulated annealing-omit map for the loop6 of room temperature structure is shown in wheat and contoured at 1.0 σ level. **(c)** 2Fo-Fc simulated annealing-omit map for the loop6 of cryogenic temperature structure is shown in gray and contoured at 1.0 σ level.
